# Cysteine Mutants of the Major Facilitator Superfamily-Type Transporter CcoA Provide insight into Copper Import

**DOI:** 10.1101/2021.05.27.446088

**Authors:** Bahia Khalfaoui-Hassani, Petru-Iulian Trasnea, Stefan Steimle, Hans-Georg Koch, Fevzi Daldal

**Affiliations:** Department of Biology, University of Pennsylvania, PA 19104, USA; Université de Pau et des Pays de l’Adour, E2S UPPA, IPREM, UMR CNRS 5254, BP1155 Pau, France; Institute of Biochemistry and Molecular Biology, ZBMZ, Faculty of Medicine, Albert-Ludwigs University of Freiburg, 79104 Freiburg, Germany

**Keywords:** Copper uptake, *cbb*_3_-type cytochrome *c* oxidase, copper-binding residues, MFS-type transporters, *Rhodobacter capsulatus* CcoA, bacterial copper import

## Abstract

CcoA belongs to the widely distributed bacterial copper (Cu) importer subfamily CalT (<u>C</u>co<u>A</u>-<u>l</u>ike <u>T</u>ransporters) of the Major Facilitator Superfamily (MFS), and provides cytoplasmic Cu needed for *cbb*_3_-type cytochrome *c* oxidase (*cbb*_3_-Cox) biogenesis. Earlier studies have supported a 12 transmembrane helices (TMH) topology of CcoA with the well-conserved Met_233_xxxMet_237_ and His_261_xxxMet_265_ motifs in its TMH7 and TMH8, respectively. Of these residues, Met_233_ and His_261_ are essential for Cu uptake and *cbb*_3_-Cox production, whereas Met_237_ and Met_265_ contribute partly to these processes. CcoA also contains five Cys residues of unknown role, and remarkably, its structural models predict that three of these are exposed to the highly oxidizing periplasm. Here, we first demonstrate that elimination of both Met_237_ and Met_265_ completely abolishes Cu uptake and *cbb*_3_-Cox production, indicating that CcoA requires at least one of these two Met residues for activity. Second, using scanning mutagenesis to probe plausible metal-interacting Met, His and Cys residues of CcoA we found that the periplasm-exposed Cys_49_ located at the end of TMH2, the Cys_247_ on a surface loop between TMH7 and THM8, and the C_367_ located at the end of TMH11 are important for CcoA function. Analyses of the single and double Cys mutants revealed the occurrence of a disulfide bond in CcoA *in vivo*, possibly related to conformational changes it undergoes during Cu import as MFS-type transporter. Our overall findings suggested a model linking Cu import for *cbb*_3_-Cox biogenesis with a thiol: disulfide oxidoreduction step, advancing our understanding of the mechanisms of CcoA function.

**Importance:** Copper (Cu) is a redox-active micronutrient that is both essential and toxic. Its cellular homeostasis is critical for supporting cuproprotein maturation while avoiding excessive oxidative stress. The Cu importer CcoA is the prototype of the widespread CalT subfamily of the MFS-type transporters. Hence, understanding its molecular mechanism of function is significant. Here we show that CcoA undergoes a thiol: disulfide oxidoreduction cycle, which is important for its Cu import activity.

## Introduction

The major facilitator superfamily (MFS) is one of the largest groups of secondary active transporters that are exceptionally diverse and ubiquitous to all three kingdoms of life. MFS members selectively transport a wide range of substrates, including sugars, amino acids, peptides and antibiotics, either by utilizing the electrochemical gradient due to differential substrate concentration, or by coupling the transport of one substrate to that of another, via the so-called alternating-access mechanism (1, 2).

A typical MFS protein comprises 400 - 600 amino acids often arranged as 12 transmembrane α-helices (TMHs) in two pseudo-symmetrical six N-terminal (N-ter) and six C-terminal (C-ter) TMH bundles, with both termini on the inner (*n*) side of the cytoplasmic membrane. In most cases, MFS proteins contain a substrate-binding cavity enclosed by the N- and C-ter domains and located about halfway into the membrane. The three dimensional (3D) structures of several MFS transporters are available and they exhibit different conformational states, supporting an alternating-access cycle mechanism of function (3–5). Accordingly, MFS-type transporters undergo a series of conformational changes to upload, and subsequently to release, its substrate across the membrane. These conformations are referred to as outward-open, outward facing-occluded, inward facing-occluded, and inward-open states. The interactions of the substrate with specific residues during its binding and its release are thought to trigger the dynamics of inter-domain interactions (3, 6). The nature and location of the residues that selectively bind the substrate and those that trigger the conformational changes required for transport may differ among the MFS proteins. In the case of the lactose permease LacY, the residues TMH4-Glu_126_ and TMH5-Arg_144_ are essential for sugar binding, whereas the TMH7-Tyr_236_, TMH8-Glu_269_ and TMH10-His_322_, which are close to one another in the inward-facing structure, coordinate the closing and opening of the protein upon substrate binding and release (7, 8). For the oligopeptide transporter PepT, the residues TMH1-Tyr_29_, TMH1-Tyr_30_ and TMH2-Tyr_68_ are important for peptide binding, while TMH10-Gly_407_ and TMH11-Trp_427_ form the pivotal points that control the occluded and the inward-facing conformational states of PepT (9).

The MFS-type transporter CcoA was the first copper (Cu) importer reported in bacteria (10-12), and has become the prototype of the large CalT (<u>C</u>co<u>A</u>-<u>l</u>ike <u>T</u>ransporters) subfamily of Cu transporters (13). It was initially identified in the facultative phototroph *Rhodobacter capsulatus* where it is required for the biogenesis of the binuclear heme-Cu (Cu_B_) center of *cbb*_3_-type cytochrome *c* oxidases (*cbb*_3_-Cox) (14), a C-type heme-Cu: O_2_ reductase. Comparative phylogenomics of CcoA orthologs showed that they are widespread among the α-proteobacteria (15). In species like *Rhodobacter sphaeroides*, which produces multiple heme-Cu: O_2_ reductases, CcoA is specific to *cbb*_3_-Cox and not involved in the maturation of the closely related *aa*_3_-type Cox (15). This is noteworthy because both the *cbb*_3_-Cox and *aa*_3_-Cox have similar heme-Cu_B_ centers (16). Distant orthologs of *R. capsulatus* CcoA, initially thought to transport riboflavin, were shown to import Cu (13), and their occurrence in some proteobacteria lacking *cbb*_3_-Cox suggests that CcoA-imported Cu is likely destined to other cuproproteins in these species. Thus, the CalT family members might have a broader role extending beyond the *cbb*_3_-Cox biogenesis.

Previous studies addressing the Cu binding and import functions of CcoA revealed two motifs, M_233_xxxM_237_ and H_261_xxxM_265_ that are well conserved among its homologs (17). The membrane topology of CcoA and locations of these motifs in the predicted TMH7 and TMH8 suggested that they are parts of its membrane-buried Cu binding site (**Fig. 1**, left panel). Substitution of these residues with alanine, which is unable to ligand metals, and analyses of the ensuing mutants for ^64^Cu uptake and *cbb*_3_-Cox production had shown that the M_233_ and H_261_ residues are essential for CcoA activity, whereas substitution of M_237_, or M_265_, which are also parts of the conserved motifs, only partially affected the function (17). The putative Cu binding site of CcoA suggested that its mode of action was likely to be different from other well-studied eukaryotic Cu transporters, such as the Ctr-type (18) or the P-type ATPase proteins (19).

**Figure 1:**
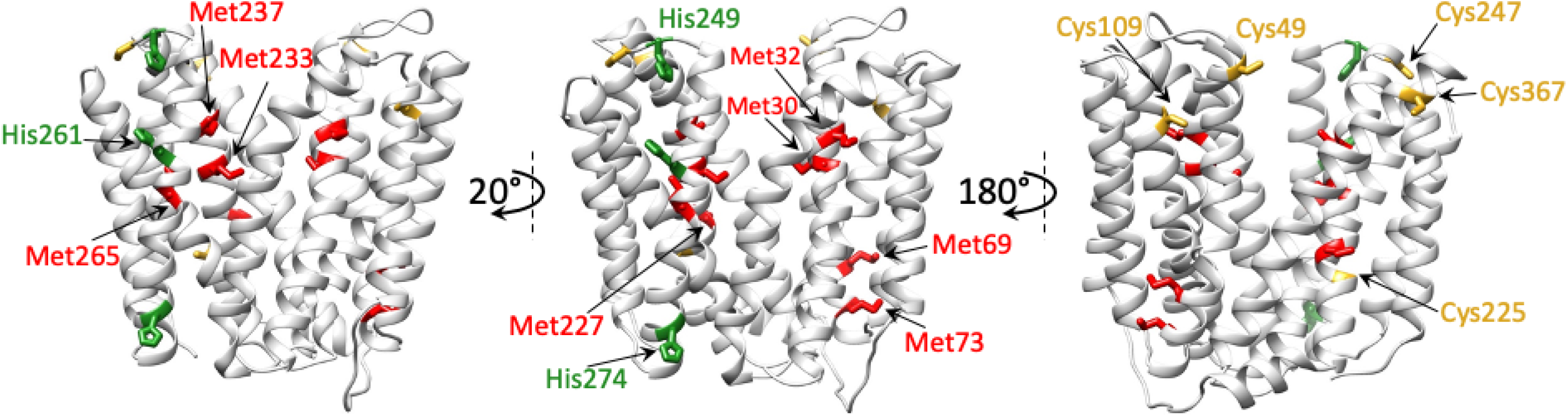
CcoA homology model using YajR as a template (CcoA_YajR_) and locations of its selected Met, His and Cys residues. Different views of CcoA_YajR_ model generated using http://swissmodel.expasy.org (GMQE: 0.51) were shown to highlight the respective locations of the conserved motifs (M_233_xxxM_237_ and H_261_xxxM_265_) proposed to bind Cu (left), the selected Met (M_30_, M_32_, M_69_, M_73_ and Met_227_) and His (H_249_ and H_274_) (middle), and the Cys (C_49_, C_109_, C_225_, C_247_ and C_367_) residues of CcoA (right).The Met, His and Cys residues are colored in red, green and yellow, respectively.

In this work, Cu import by *R. capsulatus* CcoA was studied in further detail. First, to better define the role of the M_237_ and M_265_ residues, a double mutant M_237_A+M_265_A was obtained and shown to be completely defective for Cu import and *cbb*_3_-Cox activity, in contrast to the respective single mutants. Additionally, the roles of selected five Met (M_30_, M_32_, M_69_, M_73_ and M_227_), two His (H_249_ and H_274_) and five Cys (C_49_, C_109_, C_225_, C_247_ and C_367_) residues of CcoA (**Fig. 1**, middle and right panels), were examined by monitoring their *cbb*_3_-Cox activities after mutagenesis. The results support a hypothetical model for *R. capsulatus* CcoA function, possibly involving thiol: disulfide exchange reactions between its periplasm-facing Cys residues during Cu import.

## RESULTS

### Either M_237_ or M_265_ residues of CcoA is required for Cu import

Earlier studies showed that mutants lacking CcoA were unable to accumulate ^64^Cu in a CcoA-specific (*i.e*., temperature-dependent) manner (12). Indeed, mutagenesis of the M_233_ and His_261_ residues of CcoA conserved motifs (M_233_XXXM_237_ and H_261_XXXM_265_) (**Fig. 1**, left panel) completely abolished *cbb*_3_-Cox activity (**Table 1**), and cellular ^64^Cu accumulation, while mutating M_237_ or M_265_ only partially decreased these activities (17). The results indicated that the conserved M_233_ and H_261_ residues of CcoA are essential for its function, likely forming its intra-membrane Cu binding site. However, this study was less informative about the role(s) of the remaining M_237_ and M_265_ residues of the CcoA conserved motifs (17). To further pursue this issue, a double mutant (M_237_A+M_265_A) lacking both of these Met residues was obtained, and its properties were compared with the corresponding single mutants. Both *E. coli* (**Fig. 2A**) and *R. capsulatus* (**Fig. 2B**) cells harboring the double mutant M_237_A+M_265_A produced CcoA variant proteins at wild type levels. The direct effects of these mutations on CcoA-dependent Cu uptake were determined by monitoring radioactive ^64^Cu accumulation in whole cells (Materials and Methods). Both *E. co*li and *R. capsulatus* cells producing this CcoA variant were deficient for ^64^Cu uptake (**Fig. 3A** and **3B**), similar to those mutants lacking CcoA. The *cbb*_3_-Cox activity of the double mutant was also very low (~2% of wild type), in contrast to ~73% and ~35% of the single M_237_A and M_265_A mutants, respectively (**Table 1**). The *R. capsulatus* strain lacking a chromosomal copy of *ccoA* and complemented with a plasmid-borne wild type allele (*∆ccoA*/plasmid-born *ccoA*) (Supplementary Information, **Table S1**) was used as a control and exhibited a *cbb*_3_-Cox activity of 846 ± 32 μmoles of TMPD oxidized/min/mg of total membrane proteins (referred to 100% in **Table 1**). Considering that the CcoA variant lacking both M_237_ and M_265_ residues was unable to import Cu and produce active *cbb*_3_-Cox, we concluded that at least one additional Met residue (preferentially M_265_, suggested by its more severe phenotype) located three residues apart from the M_233_ or H_261_ on TMH7 or TMH8, respectively is required for Cu import, probably as a Cu binding ligand.

**Table 1.**
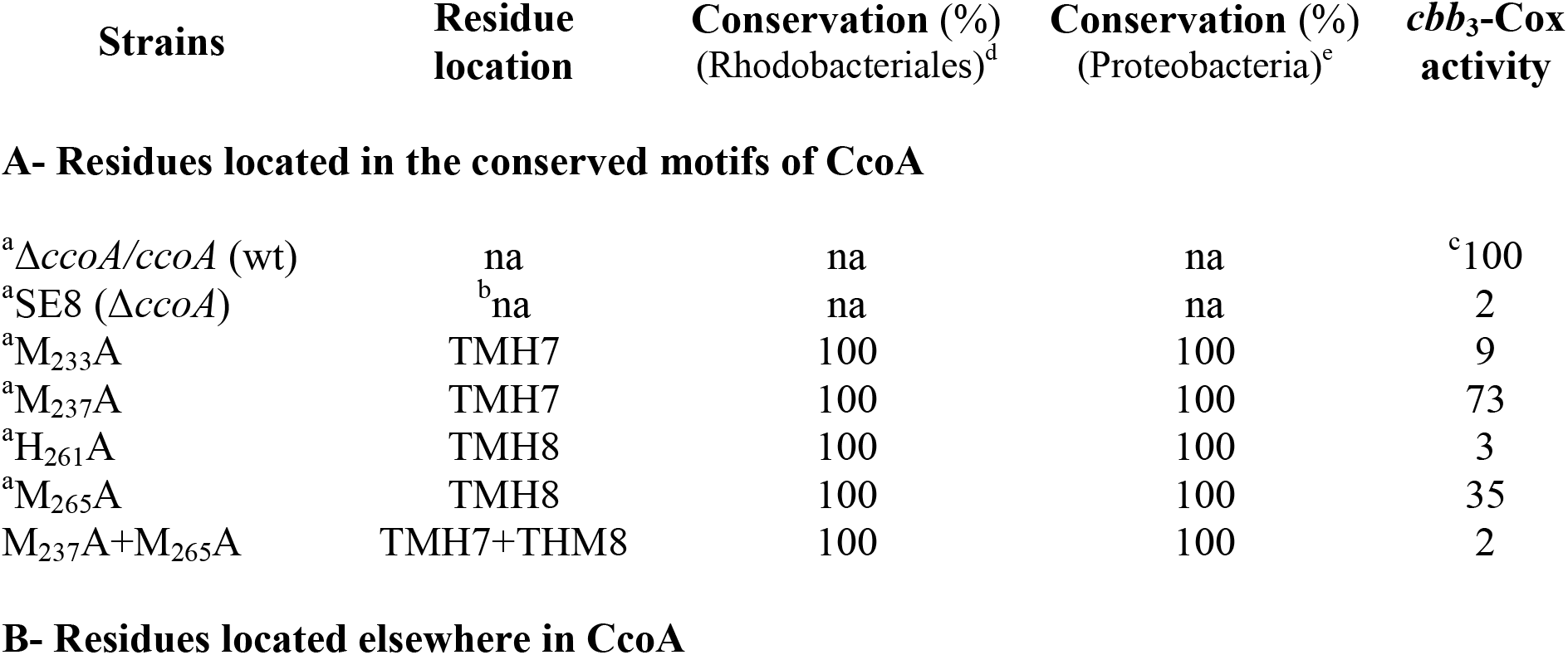
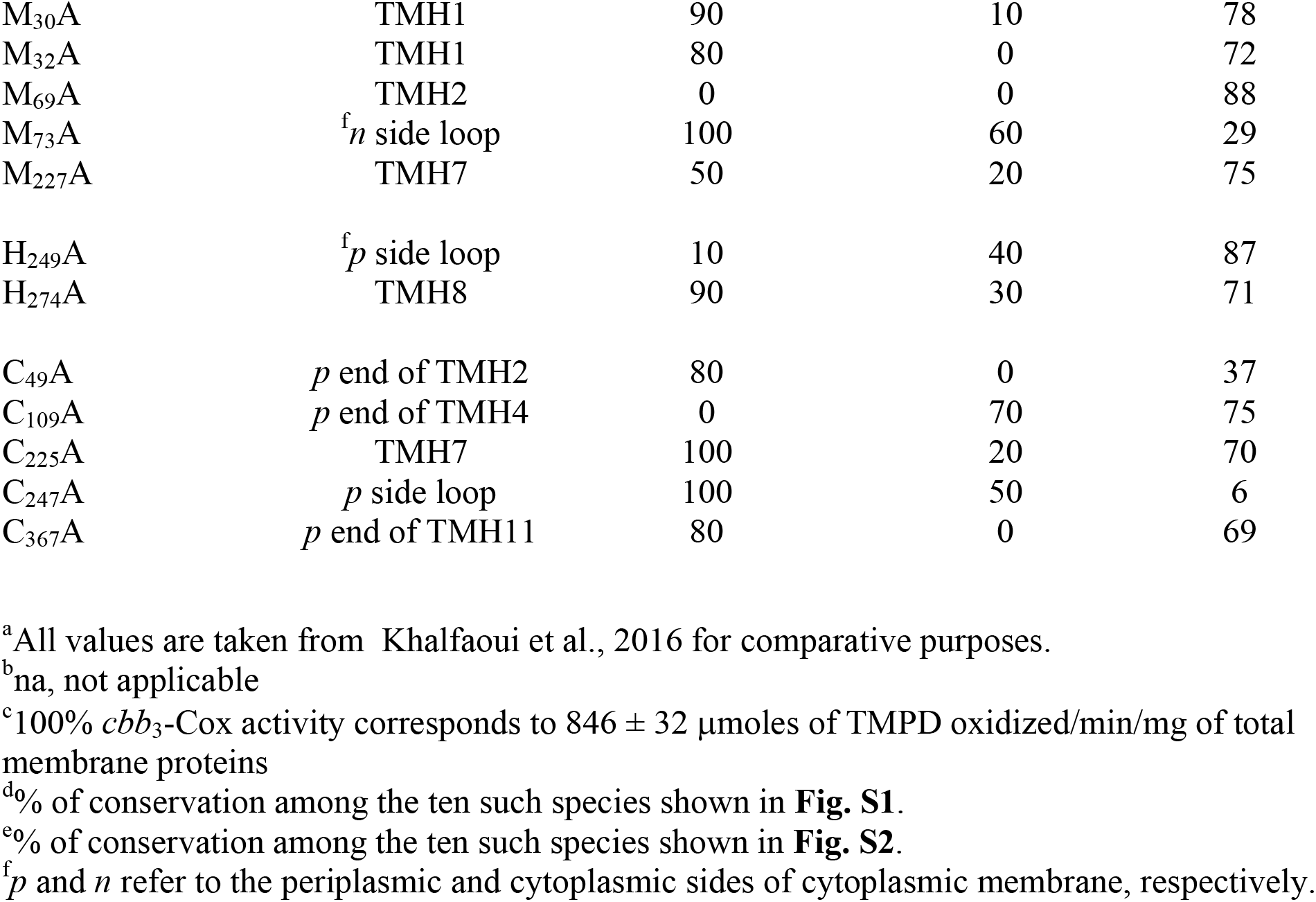
Various *R. capsulatus* CcoA mutants and their general properties.

| Strains | Residue location | Conservation (%)<br>(Rhodobacteriales)<sup>d</sup> | Conservation (%)<br>(Proteobacteria)<sup>e</sup> | *cbb<sub>3</sub>*-Cox activity |
| --- | --- | --- | --- | --- |
| **A- Residues located in the conserved motifs of CcoA** |  |  |  |  |
| <sup>a</sup>$\Delta ccoA/ccoA$ (wt) | na | na | na | <sup>c</sup>100 |
| <sup>a</sup>SE8 ($\Delta ccoA$) | <sup>b</sup>na | na | na | 2 |
| <sup>a</sup>M<sub>233</sub>A | TMH7 | 100 | 100 | 9 |
| <sup>a</sup>M<sub>237</sub>A | TMH7 | 100 | 100 | 73 |
| <sup>a</sup>H<sub>261</sub>A | TMH8 | 100 | 100 | 3 |
| <sup>a</sup>M<sub>265</sub>A | TMH8 | 100 | 100 | 35 |
| M<sub>237</sub>A+M<sub>265</sub>A | TMH7+THM8 | 100 | 100 | 2 |
| M<sub>30</sub>A | TMH1 | 90 | 10 | 78 |
| M<sub>32</sub>A | TMH1 | 80 | 0 | 72 |
| M<sub>69</sub>A | TMH2 | 0 | 0 | 88 |
| M<sub>73</sub>A | <sup>f</sup>*n* side loop | 100 | 60 | 29 |
| M<sub>227</sub>A | TMH7 | 50 | 20 | 75 |
| H<sub>249</sub>A | <sup>f</sup>*p* side loop | 10 | 40 | 87 |
| H<sub>274</sub>A | TMH8 | 90 | 30 | 71 |
| C<sub>49</sub>A | *p* end of TMH2 | 80 | 0 | 37 |
| C<sub>109</sub>A | *p* end of TMH4 | 0 | 70 | 75 |
| C<sub>225</sub>A | TMH7 | 100 | 20 | 70 |
| C<sub>247</sub>A | *p* side loop | 100 | 50 | 6 |
| C<sub>367</sub>A | *p* end of TMH11 | 80 | 0 | 69 |

<sup>a</sup>All values are taken from Khalfaoui et al., 2016 for comparative purposes.
<sup>b</sup>na, not applicable
<sup>c</sup>100% *cbb*<sub>3</sub>-Cox activity corresponds to 846 ± 32 μmoles of TMPD oxidized/min/mg of total membrane proteins
<sup>d</sup>% of conservation among the ten such species shown in **Fig. S1**.
<sup>e</sup>% of conservation among the ten such species shown in **Fig. S2**.
<sup>f</sup>*p* and *n* refer to the periplasmic and cytoplasmic sides of cytoplasmic membrane, respectively.

**Figure 2.**
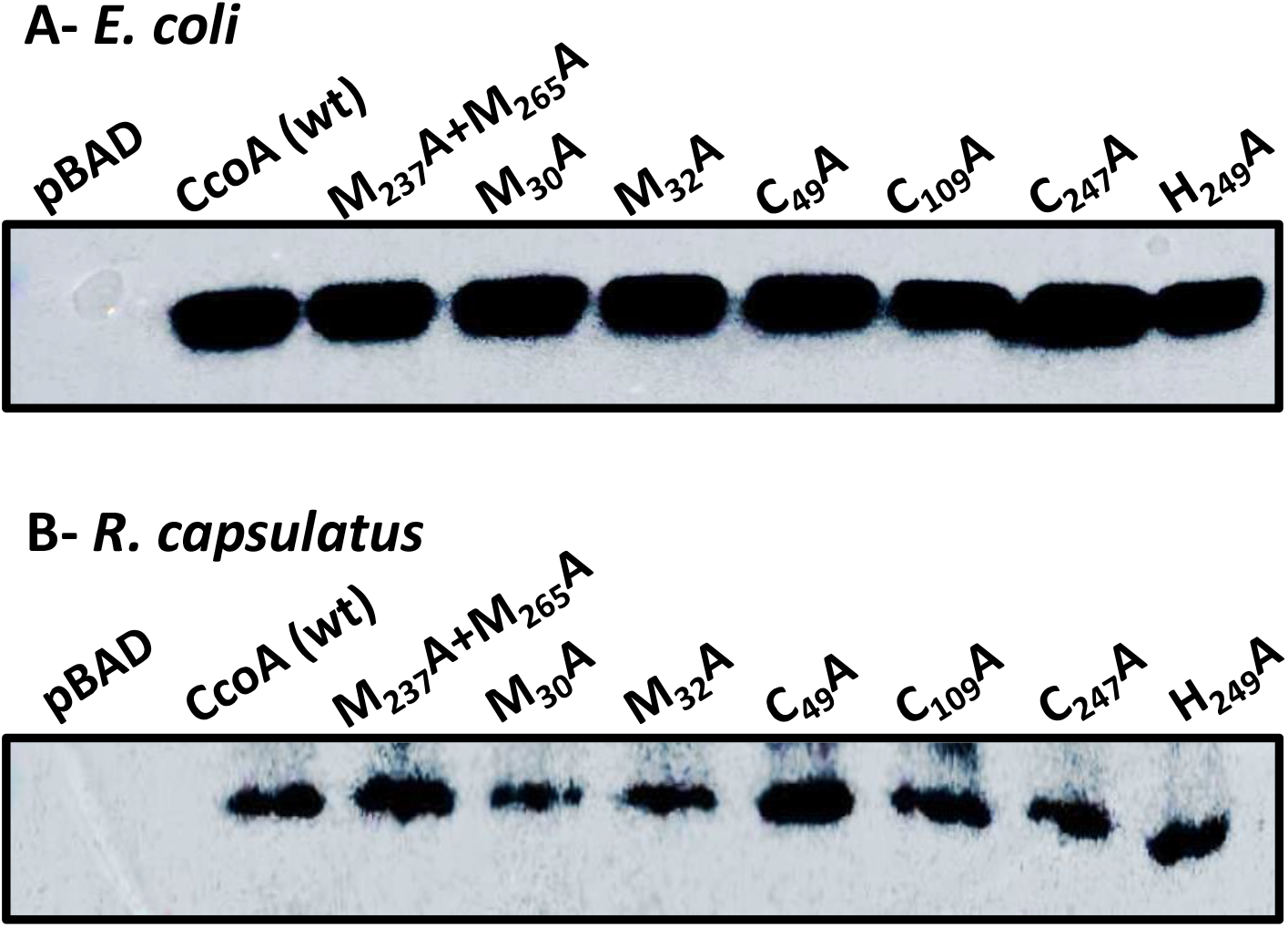
Production of mutant CcoA variants in *E. coli* and *R. capsulatus*. Membranes prepared from *L-*ara induced *E. coli* (LMG194) (**A**) and *R. capsulatus* Δ*ccoA* (SE8) **(B**) strains harboring appropriate plasmids (**Table S1**) expressing wild type or indicated CcoA mutant variants were probed with anti-Myc monoclonal antibodies. pBAD and CcoA correspond to membranes prepared from *E. coli* (**A**) or *R. capsulatus* (**B)**strains harboring empty pBAD (*E. coli*) or pRK-pBAD (*R. capsulatus*) expression plasmids, and their derivatives containing Myc-tagged versions of wild type and mutant *ccoA* alleles, as appropriate. All native and mutant proteins were produced adequately in both backgrounds.

**Figure 3.**
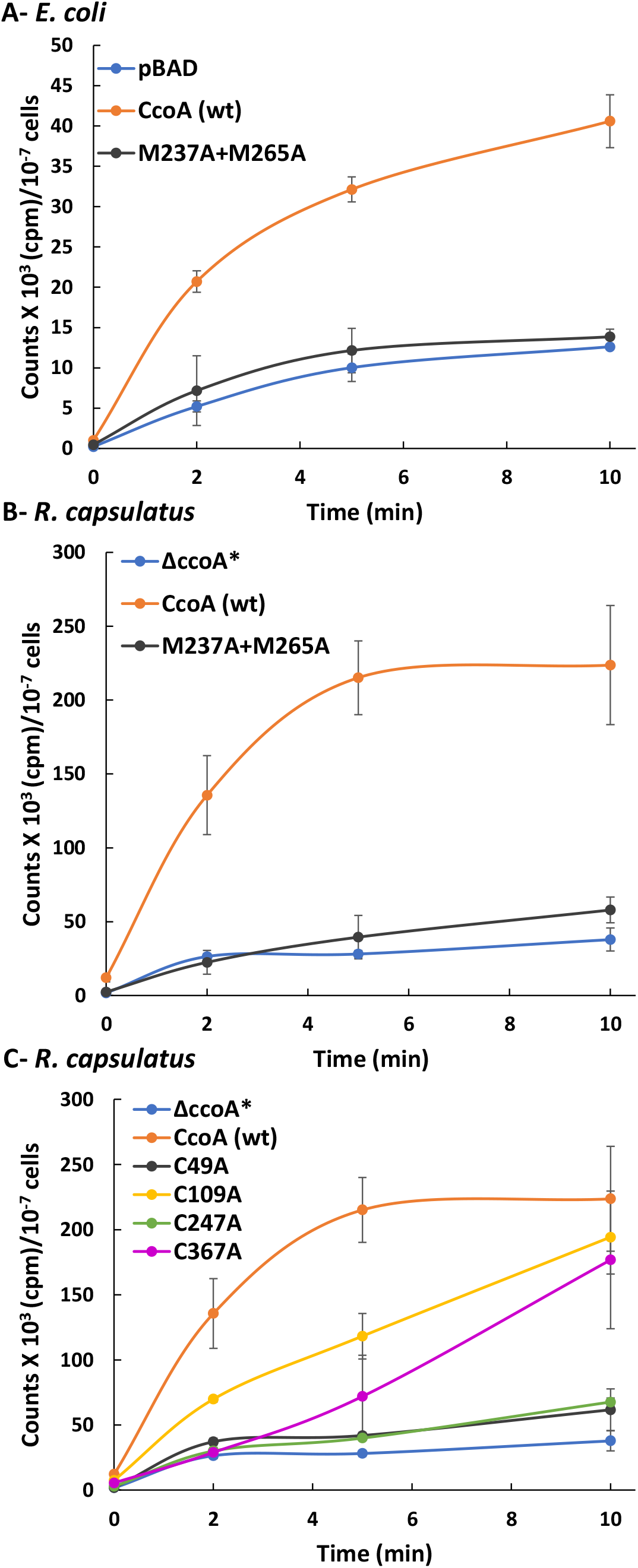
Whole cell radioactive ^64^Cu uptake by CcoA M_237_A+M_265_A double and Cys single mutant derivatives. ^64^Cu uptake kinetics observed with the CcoA M_237_A+M_265_A double mutant and C_49_A, C_109_A, C_247_A and C_367_A single mutants using appropriate *L-ara* induced (**A**) *E. coli* (LMG194) and (**B** and **C**) *R. capsulatus* Δ*ccoA ΔcopA* (SE24) cells, expressing *L-ara* inducible native and mutant *ccoA* alleles. pBAD and Δ*ccoA* refer to control strains, as appropriate. Uptake assays were performed as described in Materials and Methods. Activities measured in cells kept on ice were subtracted from those measured in cells incubated at 37 °C, and assays were repeated at least three times using multiple independent cultures. CcoA (wt) refers to (Δ*ccoA*+plasmid-borne *ccoA*) strain carrying *L-ara* inducible native CcoA, where Δ*ccoA* is Δ*ccoA* Δ*copA* (SE24), used to avoid frequent CopA revertants seen with a *ccoA* deletion (**Table S1**) (12). Error bars correspond to the standard deviations around the mean values. In each case, at least three biological and three technical repeats were done.

### Additional possible metal-liganding CcoA residues of functional importance

CcoA is rich in Met, His and Cys residues that can act as potential metal ligands (10), and those that are not parts of the conserved motifs were examined for their possible role(s). Two different amino acid sequence alignments were used with the proteobacterial homologs of CcoA that contributed to building the protein similarity network of the CalT subfamily (13). *R. capsulatus* CcoA sequence was first aligned with its closely related CcoA homologs among the Rhodobacterales (mostly from Node 1 in (13)) (Supplementary data, **Fig. S1**). This comparison included *R. sphaeroides* CcoA known to provide Cu to *cbb*_3_-Cox (16). The *R. capsulatus* CcoA sequence was also aligned with its more distant homologs among the different proteobacterial orders, including Rhizobiales, Burkhorderiales, Pseudomonales, Rhodospirales, Vibrionales, Oceanospiralles, Synecchocales, Alteromonadales, and Chromatiales (**Fig. S2**). This group included *Ochrobactrum anthropi* (Rhizobiales) CalT (CcoA ortholog) shown to transport Cu (13), and possibly required for the maturation of cuproproteins distinct from *cbb*_3_-Cox. Based on sequence alignments and topological locations (*i.e*., TMH or loop, **Table 1**) five Met (M_30_, M_32_, M_69_, M_73_ and M_227_), possibly forming the M_30_xM_32_ and M_69_xxxM_73_ motifs (of which M_30_ and M_73_ are better conserved), two His (H_249_ and H_274_), and five Cys (C_49_, C_109_, C_225_, C_249_ and C_367_) residues were retained for this study (**Fig. 1**, middle and right panels). They were substituted with Ala using *L-*ara inducible *ccoA*, and the mutants obtained were introduced into appropriate *E. coli* and *R. capsulatus* strains (Materials and Methods, **Table S1**). Their *cbb*_3_-Cox and ^64^Cu uptake activities were determined analogously to the conserved motifs residues (17).

### Properties of CcoA Met and His Mutants

The CcoA single Met (M_30_A, M_32_A, M_69_A, M_73_A and M_227_A) mutants were produced in amounts similar to the wild type and double mutant M_237_A+M_265_A in both *E. coli* and *R. capsulatus* strains (**Fig. 2**, as the data are identical for all mutants tested, only M_30_A and M_32_A are shown). Of these residues, the M_30_, M_32_, M_69_ and M_73_ form the socalled “Met (MxM and MxxxM) motifs”, sometimes implicated in binding Cu(I) (20). The single M_30_A, M_32_A and M_69_A on the N-ter domain and the M_227_A single mutant located close to the M_233_xxxM_237_ putative Cu binding motif on TMH7 had *cbb*_3_-Cox activities slightly lower than that of wild type CcoA (~78%, ~72%, ~88% and ~75% of wild type amounts, respectively), similar to the single mutant M_237_A (~73%) (**Table 1**). In contrast, mutating M_73_ that is located near the cytoplasmic end of TMH2 and highly conserved among the Rhodobacteriales CcoA homologs, led to a markedly lower c*bb*_3_-Cox activity (~29%) like the single mutant M_265_A (~35%) (**Table 1**) (17). The His_249_ and H_274_ are near the *p* and *n* sides of the membrane, respectively, and only the latter residue is conserved in Rhodobacteriales (**Fig. S1**). The corresponding mutants had *cbb*_3_-Cox activities slightly lower than wild type (~87% and ~71%, respectively) (**Table 1**). Thus, unlike M_233_ and H_261_, none of these Met and His residues were required for CcoA function, except M_73_ located close to the cytoplasm, as its substitution significantly reduces *cbb*_3_-Cox activity.

### Properties of CcoA Cys Mutants

Of the five Cys residues of CcoA, four (except C_109_) are well-conserved especially among the Rhodobacteriales (**Fig. S1**). Based on the homology model of CcoA (CcoA_YajR_) obtained using its highest homolog *E. coli* YajR (PDB: 3WDO) as a template, the C_49_ and C_367_ residues are at or near the periplasmic ends of the TMH2 and TMH11, respectively, while C_247_ is located on a periplasmic loop between TMH7 and TMH8 (**Fig. 1**, left panel) (10, 17). The non-conserved C_109_ on TMH3, and the partly conserved C_225_ on TMH7 are further embedded into the membrane, closer to the *p* and *n* sides, respectively. Substitution of each of these Cys residues with Ala did not affect the production of mutant proteins either in *E. coli* or *R. capsulatus* (**Fig. 2**, only C_49_, C_109_ and C_247_ are shown). However it impaired the *cbb*_3_-Cox activity of the mutant strains to different extents (**Table 1**). While the effects of C_109_A, C_225_A and C_367_A mutations were milder (~75%, ~70%%, and ~69% of wild type activity, respectively), those of the C_49_A and C_247_A were more severe (~37% and ~6%, respectively). In respect to Cu import, upon *L*-ara addition radioactive ^64^Cu accumulation in whole cells of a control strain lacking CcoA but harboring a plasmid-borne inducible CcoA (*∆ccoA*+plasmid-borne *ccoA*) steadily increased, unlike the *∆ccoA* strain lacking CcoA. With the C_49_A or C_247_A mutants ^64^Cu accumulation was very low, similar to the strain lacking CcoA, indicating that these residues are critical for CcoA function (**Fig. 3C**). On the other hand, the C_109_A and C_367_A mutants accumulated ^64^Cu markedly more slowly to a slightly lower level than the control cells, indicating that C_109_ and C_367_ also contribute to Cu uptake but are not essential. Overall data indicated that these mutants fall into two groups (C_49_A and C_247_A *versus* C_109_A and C_367_A) with distinct kinetics behaviors suggesting likely different functions.

### Topological locations of periplasm-facing Cys residues of CcoA

Currently, no three-dimensional (3D) structure of CcoA is available beyond the CcoA_YajR_ homology model (Global Model Quality Estimate, GMQE: 0.51) based on its most pronounced homolog which is the *E. coli* YajR (PDB:3WDO) (17). Fortunately, additional homology models of CcoA of similar GMQE yielding similar outcomes can be generated using available X-ray structures, including the iron exporter BpFPN (21). Here, we opted for two of its importer homologs, LacY (PDB:6C9W, CcoA_LacY,_ GMQE: 0.46) and GlpT (PDB:1PW4, CcoA_GlpT_, GMQE: 0.44) captured in different conformations than CcoA_YajR_. While an outward-open conformation (*i.e*., ready to receive the substrate from the *p* side of the membrane) is seen with CcoA_YajR_, the CcoA_LacY_ and CcoA_GlpT_ models provide the inward-occluded and inward-open (*i.e*., ready to release the substrate to the *n* side of the membrane) conformations, respectively (**Fig. 4A**). Top views of these models clearly show that the distances separating the periplasm facing Cys residues change drastically depending on the conformations (**Fig. 4B**) **(Table S3**, lists all appropriate α-C to α-C distances). When CcoA_YajR_ is in the outward-open conformation (**Fig. 4B** top), C_49_ and C_109_ located on the N-ter domain are very close to each other (C_49_- C_109_, 12Å), and distant from C_247_ (C_49_ - C_247_, 32Å and C_109_ - C_247_, 39Å) and C_367_ (C_49_ - C_367_, 22Å and C_109_ - C_367_, 32Å) located on the C-ter domain. In the occluded conformation of CcoA_LacY_ (**Fig. 4B** middle), C_49_ moves closer to C_247_ (C_49_ - C_247_, 23Å) and to C_367_ (C_49_ - C_367_,16Å), while C_109_ shifts closer to both C_247_ and C_367_ (C_109_ - C_247_, 36Å and C_109_ - C_367_, 27Å). In the inward-open conformation of CcoA_GlpT_ (**Fig. 4B** bottom), C_49_ and C_109_ approach even closer to C_247_ (C_49_ - C_247_, 16Å and C_109_ - C_247_, 27Å) and to C_367_ (C_49_ - C_367_, 10Å and C_109_ - C_367_, 22Å). In all conformations, the N-ter located C_49_ - C_109_ pair stays within 12-14Å, and the C-ter located C_247_ - C_367_ pair remain within 19-22Å of each other. Indeed, these distance estimations are approximations in the absence of 3D structures. Nonetheless, they depict the progressive movement of the N-ter domain C_49_ towards the C-ter domain C_247_ - C_367_ pair during the transition from the outward-open to the inward-open conformations. This observation enticed us to inquire whether the predicted distance changes between these Cys residues that are exposed to the oxidizing periplasm could undergo thiol:disulfide oxidoreduction during CcoA function.

**Figure 4.**
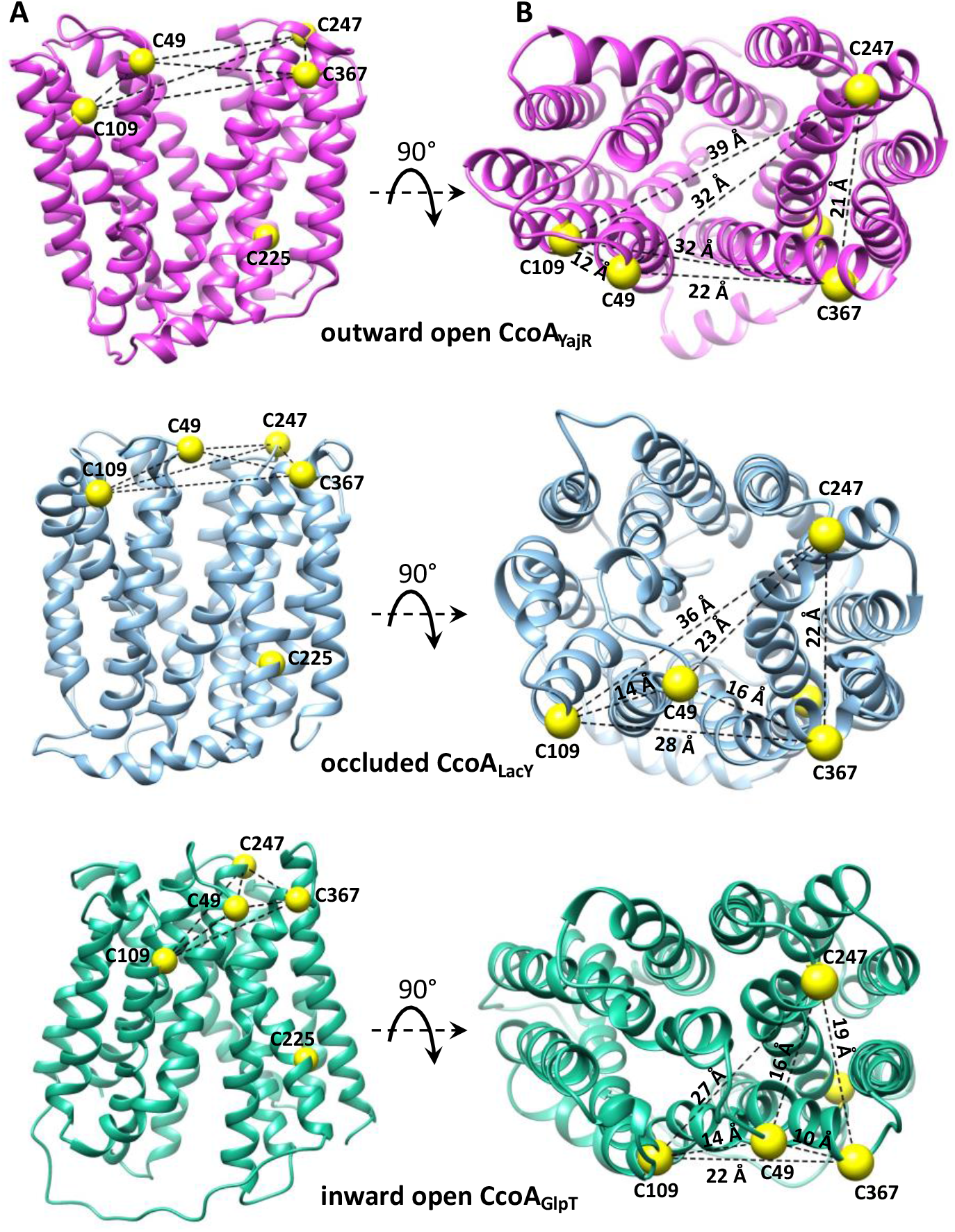
Different CcoA conformations and the distances that separate the periplasmic Cys residues in each conformational state. **A.** Side views of *R. capsulatus* CcoA homology models (CcoA_YajR_, CcoA_LacY_ and CcoA_GlpT_) representing its different conformations using as templates *E. coli* YaJR (3WDO) in the outward-facing state (16% identity; GMQE = 0.51), lactose permease LacY (1PV6) in the occluded state (12.7% identity; GMQE = 0.41) and GlpT (1PW4) in the inward-facing state (11.7% identity; GMQE = 0.46) (http://swissmodel.expasy.org). **B.** Top views of the homology models shown in **A**, with the distances separating the periplasm-facing Cys residues (yellow balls) in different conformations of CcoA.

### The disulfide bonds formed between the Cys residues of CcoA

The occurrence *in vivo* of disulfide bond(s) in CcoA was probed using *E. coli* cells expressing native CcoA or its Cys mutant variants and the thiol-reactive alkylating agent monomethyl-(PEG)24-Maleimide (mPEG) (22). Alkylation of free Cys thiols of CcoA by mPEG is expected to increase its MW by ~1.2 kDa per free thiol. In the case of disulfide bonds, alkylation occurs only after reduction by DTT, and then mPEG further increases the MW by ~2.4 kDa per reduced disulfide bond. The relative MW changes (Mr) in native and Cys mutant variants of CcoA were followed by SDS-PAGE/immunodetection (**Figs. 5** and **6**).

**Figure 5:**
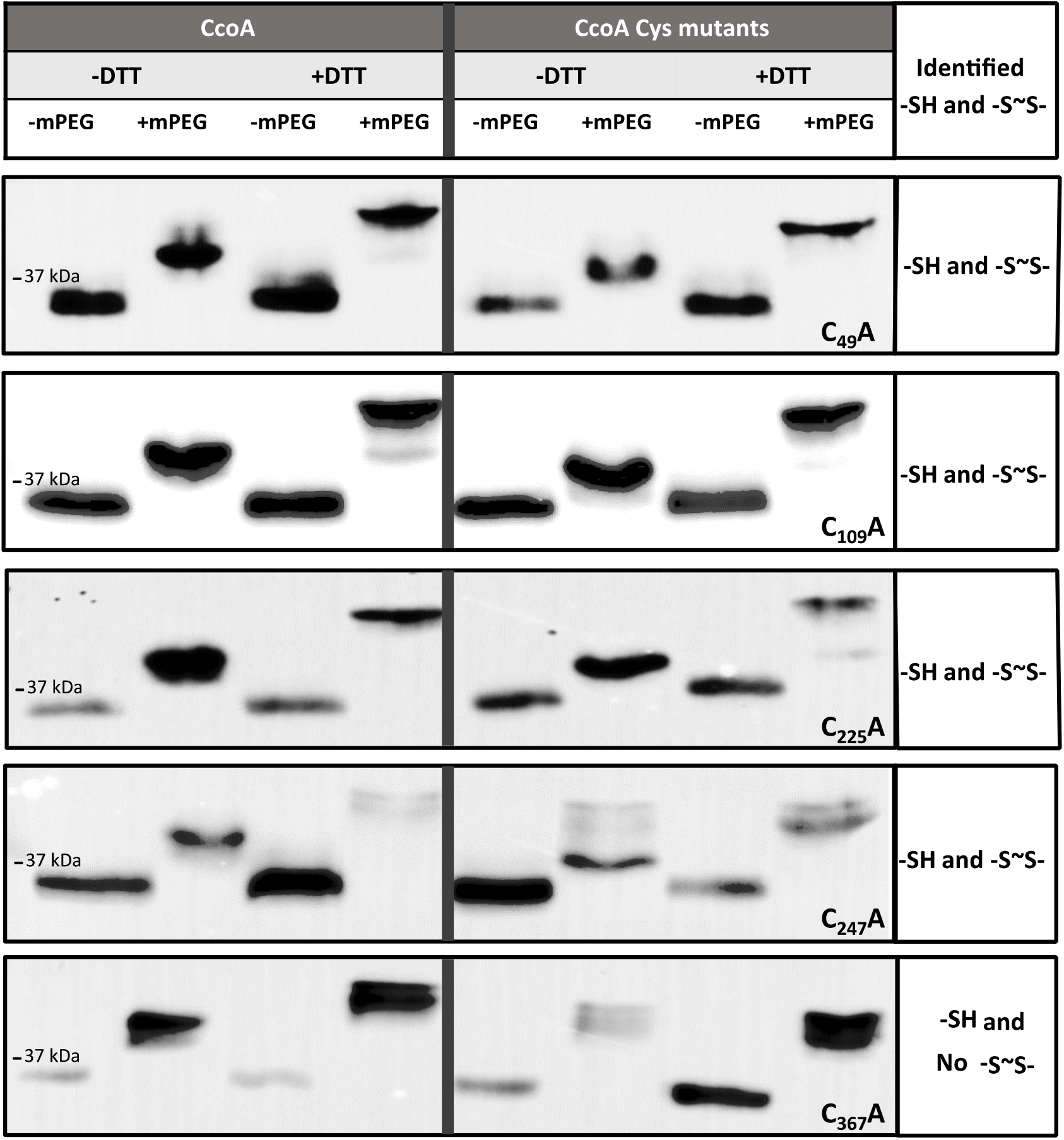
*In vivo* free thiols and disulfide bonds in native CcoA and its single Cys mutant derivatives. SDS-PAGE followed by immunoblot analysis of *E. coli* cells expressing native and single C_49_A, C_109_A, C_225_A, C_247_A and C_367_A mutant variants of CcoA. Cells growing in the presence of *L-ara* were treated mid-log phase for 10 min with or without DTT, followed by TCA precipitation and subsequent alkylation with or without mPEG. The presence of unmodified or mPEG alkylated CcoA was detected using anti-Myc monoclonal antibody and horseradish peroxidase conjugated anti-mouse IgG (Materials and Methods). Total protein amounts of SDS solubilized TCA-precipitated pellets could not be determined precisely, leading to variations of protein amounts loaded per lane. Thus, only the qualitive occurrence of Mr shifts in the absence or presence of DTT or mPEG were taken into consideration in this experiment.

**Figure 6:**
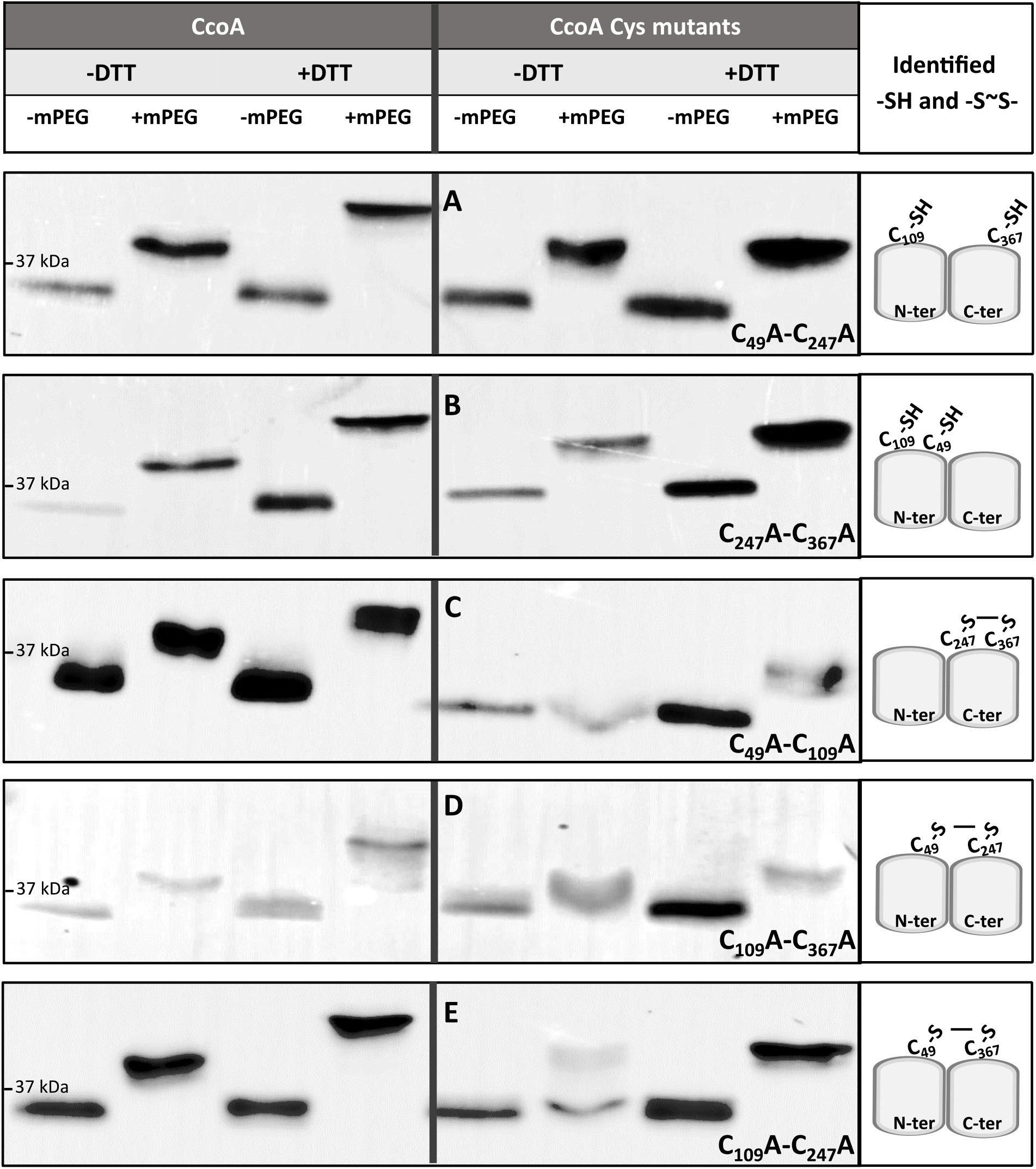
*In vivo* free thiols and disulfide bonds formed in native CcoA and its double Cys mutant derivatives. The experiments were conducted as described in Fig. 5, except that *E. coli* cells expressing the native (left side panels) and double mutant variants of CcoA were used: (**A**) C_49_A - C_247_A; (**B**) C_247_A - C_367_A; (**C**) C_49_A - C_109_A; (**D**) C_109_A - C_367_A; (**E**) C_109_A - C_247_A. As in Fig. 5, total protein amounts of SDS solubilized TCA-precipitated pellets could not be determined precisely, leading to variations of protein amounts loaded per lane. Thus, only the Mr shifts detected in the absence or presence of DTT or mPEG were considered in these experiments.

Under our conditions, native CcoA (predicted MW of 37.4 kDa) runs as a band of ~35 kDa Mr in its oxidized or reduced forms (**Fig. 5**, left panels), which is not uncommon for membrane proteins. In the absence of DTT, alkylating by mPEG increased native CcoA Mr by ~3-4 kDa to ~38-39 kDa, suggesting that it contained three free Cys thiols (predicted Mr of 38.6 kDa after three mPEG addition). Moreover, alkylating native CcoA after DTT reduction further increased its Mr by another ~2-3 kDa to ~ 40-41 kDa, indicating that the native protein contained one disulfide bound *in vivo* (**Fig. 5**, left panels). Although detecting the ~1-2 kDa Mr differences with hydrophobic membrane proteins was challenging, following TCA precipitation all CcoA Cys residues appeared accessible to alkylation, including the Cys_109_ and Cys_225_ which are more buried into the lipid bilayer according to the CcoA structural models (**Figs. 1** and **4**).

Similar mPEG alkylation experiments were repeated using single Cys mutants variants of CcoA (**Fig. 5**, right panels). Without DTT reduction, all single Cys mutant variants exhibited mPEG-induced Mr shifts similar to native CcoA, and the largest shift was seen with C_367_A mutant. In all cases but C_367_A, the shifts were consistent with the likely presence of at least two free thiols, but not four as it would have been expected upon elimination of any Cys residues already engaged in a disulfide bond in native CcoA. Following DTT reduction, all single Cys mutants, except C_367_A, showed the additional mPEG-induced Mr shifts, indicating that they still contained a disulfide bond formed among the remaining Cys residues. This observation suggested that native CcoA has more than two Cys residues that could form a disulfide bond(s). The C_367_A mutant was intensely alkylated but did not exhibit any readily detected mPEG-mediated Mr shift after DTT reduction, indicating that it contained no more disulfide bonds, and suggested that this residue provides one of the thiol groups forming a stable disulfide bond in native CcoA (**Fig. 5**, right panel, last row). Although occasionally additional minor bands were also seen in some cases (*e.g*., native CcoA, **Fig. 5** left panel second row, or C_225_A, **Fig. 5** right panel third row), the data showed that the C_49_A, C_109_A, C_225_A or C_247_A single Cys mutants behaved similarly to each other and to native CcoA, which precluded identification of partner cysteines for forming a disulfide bond.

To identify the disulfide bond forming partner(s) in native CcoA *in vivo*, a set of double Cys mutants were examined (**Fig. 6**). All CcoA double Cys variants were produced adequately in *E. coli* and in *R. capsulatus*, and exhibited low *cbb*_3_-Cox activities like their cognate single Cys mutants. In the absence of DTT, mPEG alkylation data showed that the double mutants C_49_A - C_247_A (with C_109_, C_225_ and C_367_ intact) and C_247_A - C_367_A (with C_49_, C_109_ and C_225_ intact) had Mr shifts similar to each other, and to native CcoA, containing free thiols (**Fig. 6**, panels **A** and **B**).

However, like the C_367_ single mutant (**Fig. 5**, bottom row), these two double Cys mutants did not exhibit any additional mPEG-induced Mr increase upon reduction by DTT, indicating that the remaining Cys residues did not form disulfide bonds. Conversely, the double mutants C_49_- C_109_ (with C_225_, C_247_ and C_367_ intact) and C_109_- C_367_ (with C_49_, C_225_ and C_247_ intact) showed no or slight mPEG-induced Mr shifts in the absence of DTT (although the absence of this shift was less clear in the latter double mutant), but exhibited more clear Mr shifts upon mPEG alkylation after DTT treatment (**Fig. 6**, panels **C** and **D**). Since the C_225_ residue is near the *n* side and remote from the other periplasm-exposed Cys residues on the *p* side of the membrane, it is likely that in the C_49_A - C_109_A and C_109_A - C_367_A double mutants, the C_247_ and C_367_ and the C_49_ and C_247_ residues, respectively formed disulfide bonds (although the latter pair might form a less stable disulfide bond) (**Fig. 6**, panel **C** and **D**, far right). The slight Mr shifts seen with these double mutants in the absence of DTT reduction were consistent with the poor alkylation of C_225_, still intact in these mutants. The remaining C_109_A - C_247_A double mutant (with C_225_, C_49_ and C_367_ intact) behaved essentially like the latter mutants, except that the CcoA population appeared heterogenous in the absence of DTT (**Fig. 6**, panel **E**). A small fraction contained free thiols that was alkylated by mPEG without DTT treatment, whereas a large fraction contained a disulfide bond that could only be alkylated after DTT reduction. Again, assuming that C_225_ is too far from the other Cys residues to participate in disulfide bond formation, a large fraction of the C_109_A - C_247_A double mutant population comprises a less stable disulfide bond between the C_49_ and C_367_ residues (**Fig. 6**, panel E). In summary, the overall data indicated the formation of disulfide bonds between C_49_~C_247_, C_49_~C_367_ and C_247_~C_367_ (with the last one forming the most stable bond), and the clear absence of disulfide bonds between C_109_~C_367_ and C_49_~C_109_. This suggests that Cys_109_ is redox-inactive, unlike the other periplasm-exposed residues, possibly due to its membrane-buried location in all conformations of CcoA (**Fig. 4**). Consequently, in cells producing native CcoA, any pair among the C_49_, C_247_ and C_367_ residues could form a single disulfide bond *in vivo*, leaving behind three free thiol groups, including C_109_ and C_225_. This finding raised the possibility that native CcoA *in vivo* might exist as a heterogenous population with different conformations, presumably due to the import of spurious Cu present in the growth medium. How the initial binding of Cu changes the conformation of CcoA, and shuffles the free thiols and disulfide bonds between its three active Cys residues, remains to be determined in future studies.

## DISCUSSION

This study focused on the role of plausible metal-liganding residues Met, His and Cys of CcoA, a member of the CalT (<u>C</u>co<u>A</u>-<u>l</u>ike <u>T</u>ransporters) subfamily of MFS-type transporters (13) and the prototype of proteobacterial Cu importers (10, 15). The CalT subfamily is characterized by two well-conserved motifs (M_233_xxxM_237_ and H_261_xxxM_265_ in *R. capsulatus* CcoA) of which the first Met and His residues are absolutely required for Cu import (17). Here we show that mutating concomitantly the M_237_ and M_265_ residues also abolishes CcoA activity, unlike the corresponding single mutants. Thus, the presence of at least one additional Met residue together with Met_233_ and His_261_ is required for Cu import. This finding further supports the Cu binding role of the conserved motifs that are the hallmark of the CalT subfamily of MFS-type transporters (13, 17).

We examined the distribution and topological location of additional possible metal-liganding residues of CcoA that are often conserved among its homologs, in particular those from the Rhodobacterales within the Proteobacteria. Of these residues, mutating C_225_, M_227_ or H_274_ located at the TM7 and TM8 on the C-ter domain of CcoA near the membrane Cu-binding site, had little effect on CcoA activity. This finding was similar to that seen with the M_237_A or M_265_A single mutants, suggesting that they were either not critical for function, or partly substituted by surrogate residues. Intriguingly, mutating M_73_, but not M_69_, of the putative “Met” motif (M_69_xxxM_73_ in *R. capsulatus*) (20) had a stronger effect on *cbb*_3_-Cox activity. Homology models of the different conformations of CcoA do not seem to suggest that these N-ter residues come very close to the C-ter Cu binding residues. However, how Cu is released from CcoA is not yet known, leaving the possibility that the C_225_, M_227_, H_274_ residues or the putative Met (M_69_xxxM_73_) motif, or both, all positioned closer to the *n* side of the membrane, might play a role in this process.

Remarkably, mutating the periplasm-exposed C_49_, C_247_ and C_367_ residues affected CcoA activity to different degrees. These residues are well-conserved among the Rhodobacterales, but either less (~50% for C_247_) or not (0% for C_49_ and C_367_) conserved in other Proteobacterial orders (**Figs. S1** and **S2**). The basis of this conservation is not obvious, but it might relate to the ultimate destination of Cu (*e.g*., *cbb*_3_-Cox in *R. capsulatus* and other cuproenzymes in *O. anthropi*) and the different Cu donors and acceptors of CcoA and its homologs. Of the periplasm-facing Cys (C_49_, C_109_, C_247_ and C_367_) residues of CcoA, mutating C_109_ slowed Cu uptake (**Fig. 3C**), slightly affected *cbb*_3_-Cox activity (**Table 1**), and mPEG alkylation indicated that C_109_ does not form a disulfide bond with either C_367_ or C_49_. Intriguingly, C_109_ residue is not conserved among the Rhodobacterales (0%), but is better conserved (~70%) among the other orders of proteobacteria where CalT is thought to provide Cu to other cuproproteins distinct from the *cbb*_3_-Cox (13).

Alkylation data of the single and double Cys mutants revealed that in native CcoA, two of the three periplasm-facing C_49_, C_247_ and C_367_ residues form a disulfide bond, while the remaining two stay as free thiol *in vivo*. Moreover, all possible disulfide and free thiol combinations among these residues (*i.e*., C_247_~C_367_ leaving C_49_ free, C_49_~C_247_ leaving C_367_ free and C_49_~C_367_ leaving C_247_ free) were observed in appropriate Cys double mutants. However, the level of stability of these bonds seem to be different, with the C_247_~C_367_ bond being most stable. Although the data in **Fig. 6** tend to suggest that the C_49_~C_247_ bond might be formed, yet the data with C_367_A mutant in **Fig. 5** suggest that it certainly must not be stable to be readily detected in this mutant. Assuming that CcoA undergoes conformational changes like any MFS-type transporter, the disulfide bond formation patterns suggest a hypothetical model linking Cu binding and conformational changes (**Fig. 7**). Accordingly, in the outward-open conformation of CcoA (**state 1**), C_247_ and C_367_ would contain a disulfide bond, far away from C_49_. Binding of Cu would convert CcoA into its occluded conformation (**state 2**), bringing C_49_ near the C_247_~C_367_ disulfide bond, and a nucleophilic attack would yield either C_49_~C_367_ (**Fig. 7**, left) or C_49_~C_247_ (**Fig. 7**, right) disulfide bond while freeing the remaining thiol of C_247_ or C_367_. We note that if no such disulfide bond is formed or is extremely unstable, then the occluded conformation (**state 2**) may not have a disulfide bond (not shown in **Fig. 7**). Exponentially growing cells used in this study not being synchronized for Cu import, different conformations of CcoA must co-exist, rendering impossible to discriminate between these possibilities at this stage. The more defective phenotype and the periplasmic location (*i.e*., increased solvent exposure) of C_247_ and the weaker nature of C_49_~C_247_ (as suggested by its absence in C_367_A single mutant) and the detection of C_49_~C_367_ (as seen with C_109_A - C_247_A double mutant) disulfide bonds might argue that the C_49_- C_367_ disulfide bond may be more favorable at the inward open conformation (**state 3**) **(Fig. 7**, right). In any case, further progression of Cu within CcoA from the periplasm towards the cytoplasm would trigger the remaining free thiol (C_247_ or C_367_) to attack the disulfide bond involving C_49_ (C_49_~C_247_ or C_49_~C_367_) at the inward open conformation (**state 3**). The subsequent resolution of this bond would then re-establish the initial C_247_~C_367_ disulfide bond and free C_49_ thiol, returning CcoA to its outward-open conformation (**state 1**). This model attributing more critical roles to C_49_ and C_247_ is also consistent with the highly defective ^64^Cu uptake seen with the C_49_A and C_247_A single mutants (**Fig. 3C**). Conceivably, the three periplasm-facing C_49_, C_247_ and C_367_ residues that are highly conserved in Rhodobacteriales may also play additional and perhaps different roles (*e.g*., liganding Cu) instead of those ascribed here. Yet, this hypothetical model suggests a link between the binding of Cu, ensuing conformation changes, and plausible thiol: disulfide oxidoreduction of CcoA. In this respect, the absence of *R. capsulatus* thiol: disulfide oxidoreductase DsbA (23), which catalyzes intramolecular disulfide bonds in extra-cytoplasmic proteins, is known to affect *cbb*_3_-Cox biogenesis (24). Whether or not DsbA is involved in these thiol: disulfide exchange reactions seen with CcoA is presently unknown, but future studies addressing determination of the thiol: disulfide exchange reaction rates (*e.g*., using 5,5-dithio-bis-2-nitrobenzoic acid) (23) and the pKa values of appropriate thiols might identify the attacking and resolving Cys residues to further elucidate this process.

**Figure 7.**
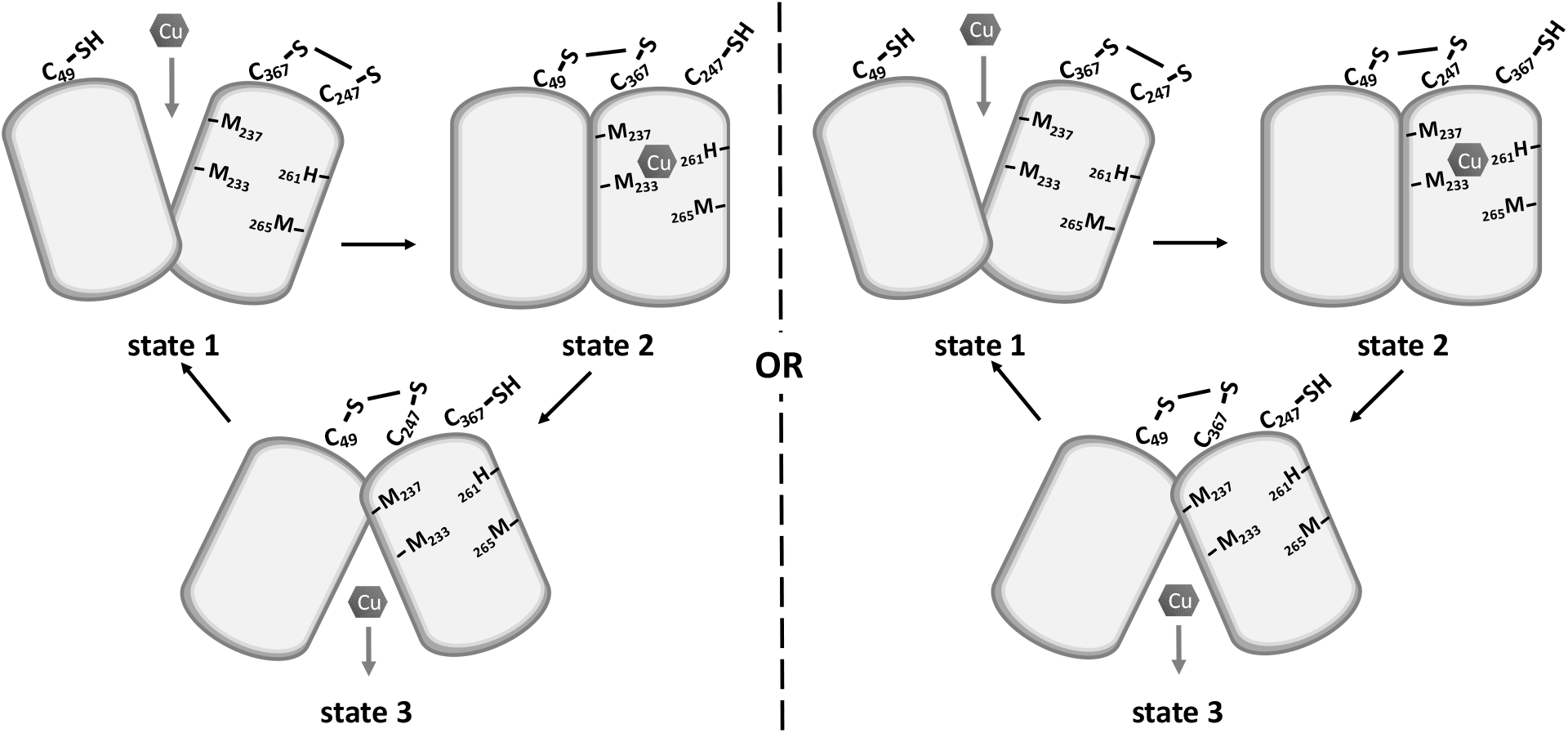
A hypothetical model linking substrate binding, conformational changes and thiol:disulfide exchange reactions that CcoA might undergo during Cu import. Accordingly, at the onset CcoA has one disulfide C_247_~C_367_ and C_49_-SH (outward open **state 1**), Binding of Cu induces a conformational change leading to occluded **state 2**where C_49_~C_367_ (left side) or C_49_~C_247_ (right side) disulfide bond is formed (although the available data cannot rule out the absence of a disulfide bond at this state). Exchange of disulfide bonds and progress of Cu from the *p* to the *n* side of the membrane yield the inward open **state 3**. Upon release of Cu and resolution of the latter disulfide bond, CcoA is returned to its starting conformation.

Noticeably, other Cu transporters also have critical Cys residues. The Ctr-type transporters contain important Cys residues (within the HCH motif) at their C-terminal parts, near the *n* side of the membrane (25). They form disulfide bonds between the monomers of trimeric CTR1 to facilitate Cu release, unlike the periplasm-facing Cys residues of CcoA monomer, presumably involved in Cu uptake. Clearly, a requirement for Met, His and Cys residues for CcoA activity distinguishes its mechanism of function from other well-known Cu(I) transporters, including the eukaryotic Ctr1 (25-27) or the bacterial CopA and CusA proteins (28, 29).

In summary, this work showed that the periplasm-facing Cys residues, together with the conserved membrane-embedded Cu-binding (M_233_xxxM_237_) and (H_261_xxxM_265_) motifs, are critical for proper function of the MFS-type Cu importer CcoA, and possibly of its close homologs among Rhodobacteriales.

## MATERIALS AND METHODS

### Growth conditions, strains and plasmids used

The bacterial strains used in this work are described in **Table S1**. *Escherichia coli* strains were grown at 37 °C on Luria Bertani (LB) medium, supplemented with antibiotics (100 and 12.5 μg/mL ampicillin (Amp) and tetracycline (Tet), respectively) and L-arabinose (*L-*ara) (0.5%), as appropriate (17). The purple non-sulfur facultative photosynthetic *R. capsulatus* strains were grown at 35 °C under respiratory (aerobic dark) conditions in enriched MPYE medium supplemented with antibiotics (2.5 μg/mL Tet) and *L*-Ara (0.5%), as needed (30).

### Construction of CcoA mutants

Standard molecular genetic techniques were performed as described by (31). The plasmids and primer sequences used are listed in **Tables S1** and **S2**, respectively. The plasmid pBK68 carrying the *L*-ara inducible *ccoA* is a derivative of pBAD/Myc-HisA (17), and used as a template for site-directed mutagenesis. Appropriate forward and reverse mutagenesis primers (**Table S2**) were used to generate the plasmids pBK98 (M_30_A), pBK99 (M_32_A), pBK100 (M_69_A), pBK101 (M_73_A), pSP6 (M_227_A), pSP4 (H_249_A), pSP5 (H_274_A), pBK108 (C_49_A), pBK109 (C_109_A), pSP9 (C_225_A), pSP8 (C_247_A), and pBK117 (C_367_A), producing CcoA variants with the indicated mutations (**Table S1**). These plasmids served as templates for generating the *ccoA* double Met or Cys mutants by using appropriate forward and reverse primers (**Table S2**) to yield pBK79 (M_237_A+M_265_A), pBK119 (C_49_A+C_109_A), pBK122 (C_49_A+C_247_A), pBK126 (C_109_A+C_247_A), pBK123 (C_247_A+C_367_A), and pBK129 (C_109_A+C_367_A), producing mutant CcoA variants (**Table S1**). The pBAD-derivatives carrying the mutant alleles of *ccoA* were cut with NsiI and ligated to the PstI site of the broad host-range plasmid pRK415 to generate the following pBAD-pRK415 composite plasmid derivatives carrying the mutant *ccoA* alleles, as follows: pBK90 (H_249_A), pBK95 (M_237_A+M_265_A), pBK102 (M_30_A), pBK103 (M_32_A), pBK104 (M_69_A), pBK105 (M_73_A), pBK92 (M_227_A), pBK91 (H_249_A), pBK111 (C_49_A), pBK112 (C_109_A), pBK94 (C_225_A), pBK93 (C_247_A), pBK120 (C_367_A), pBK121 (C_49_A+C_109_A), pBK124 (C_49_A+C_247_A), pBK127 (C_109_A+C_247_A), pBK125 (C_247_A+C_367_A), and pBK130 (C_109_A+C_367_A) (**Table S1**). These pRK derivatives were conjugated into appropriate *R. capsulatus* strains, as described earlier (17).

### Chromatophore membrane preparation, SDS-PAGE and immunodetection

Intracytoplasmic membrane vesicles (chromatophore membranes) were prepared in 20 mM Tris-HCl, pH 7.0 containing 150 mM NaCl and 1 mM phenylmethylsulfonylfluoride (PMSF) according to (30). Total protein concentrations were determined using the bicinchoninic acid assay (Sigma Inc.; procedure TPRO-562), and SDS-PAGE (12%) analyses were conducted as in (32). Prior to loading, 20 or 40 μg of proteins samples of *E. coli* or *R. capsulatus*, respectively, were solubilized by incubation at room temperature for 10 min in a loading buffer (final concentration of 62.5 mM Tris-HCl pH 6.8, 2% SDS, 2% β-mercaptoethanol and 0.01% bromophenol blue). After electrophoresis, gels were electroblotted onto Immobilon-P polyvinylidene difluoride (PVDF) membranes (Millipore Inc., MA), and probed with anti c-Myc tag monoclonal antibodies (clone 9E10 from University of Pennsylvania Cell Center). The secondary antibodies were horse radish peroxidase-conjugated anti-mouse IgGs (GE, Healthcare Inc.), and the signal was detected using Supersignal West Pico chemiluminescence substrate according to the supplier’s recommendations (Thermo Fisher Inc.).

### Determination of the free thiol groups and disulfide bonds of CcoA *in vivo*

*E. coli* cells producing wild type or appropriate Cys mutant derivatives of CcoA were analyzed by alkylating free Cys thiols with methoxy-polyethylene-glycol-maleimide (mPEG-MAL) before and after dithiothreitol (DTT) reduction according to (22). Strains producing desired CcoA variants were grown overnight at 37° C in 5 mL LB medium supplemented with appropriate antibiotics (Amp, 100 μg/mL and Tet, 10 μg/mL) with 180 rpm shaking. The next day, 100-200 μl of these cultures were sub-cultured into 10 ml fresh LB medium containing 1% *L-*ara and appropriate antibiotics at 37° C with shaking (180 rpm) until they reached an OD_600_ of 0.5. At this stage, two aliquots of 0.9 mL each were taken out and kept on ice, while the remaining culture (8.2 mL) was reduced by addition of 82 μL of 1.0 M DTT (10 mM final concentration) and further incubation for 10 min at 37° C with shaking. Two additional aliquots of 0.9 mL each were taken and placed on ice. All four samples were precipitated by addition of 100 μL of 100% ice cold TCA (final concentration 10% v/v) and incubated on ice for 30 min. Precipitated materials were centrifuged at 13,000 rpm at 4° C for 12 min, supernatants removed without disturbing the pellets, which were washed with 300 μL ice cold acetone to eliminate TCA, and re-centrifuged under the same conditions. The pellets were partially dried at 30° C for ~10 min to evaporate acetone, one untreated and one DTT treated pellets were resuspended in 30 μL PBS buffer (137 mM NaCl, 2.7 mM KCl, 10 mM Na_2_HPO_4_, 2.0 mM KH_2_PO_4_, pH 7.0) supplemented with 0.1% SDS. Similarly, the remaining one untreated and one DTT treated pellets were resuspended in 30 μL mPEG-MAL solution (20 mM mPEG-MAL dissolved in PBS buffer) supplemented with 0.1% SDS. The pellets were vortexed vigorously for 1 min for complete dissolution, and incubated in the dark, at room temperature for 2 h under constant shaking (1000 rpm) to label the accessible thiol groups of CcoA. At the end of the incubation, 10 μL of 5 x Laemmli buffer (10 % SDS (v/v), 0.05% bromophenol blue (w/v), 60% glycerol (v/v), 300 mM Tris-HCl, pH 6.8) were added to each sample, and further incubated at room temperature for 15 min. 20 μL of each sample were loaded on a 12% non-reducing SDS-PAGE gel, run at 200 V and subjected to immunoblot analyses using α-myc monoclonal antibodies (1:5000 dilution) as primary, and horse radish peroxidase conjugated anti-mouse IgGs as secondary antibodies (1:3000 dilution). Addition of mPEG-MAL, specific to free thiol groups, increases the Mr of alkylated mPEG-MAL derivatives of CcoA, with the increases being proportional to the number of free thiol groups. Comparison of untreated and DTT treated samples prior to mPEG-MAL alkylation visualized the DTT-reduced disulfide bonds of CcoA *in vivo* under the growth conditions used.

### The *cbb*_3_-Cox activity

The *in situ cbb*_*3*_-Cox activity of *R. capsulatus* colonies was assessed qualitatively using the “NADI” staining solution, which is made by mixing in 1:1 (v/v) ratio 35 mM α-<u>na</u>phthol and 30 mM *N, N, N’, N’* –<u>di</u>methyl-*p*-phenylene diamine (DMPD) dissolved in ethanol and water, respectively (33). Colonies producing *cbb*_3_-Cox stain blue, while those lacking it remain unstained. The *in vitro cbb*_3_-Cox activity was measured quantitatively using *R. capsulatus* chromatophore membranes and tetramethyl-*p*-phenylenediamine (TMPD) by monitoring spectrophotometrically in a stirred cuvette its oxidized form at 562 nm (*λ*_562_ = 11.7) at room temperature. Briefly, 10 μg of *R. capsulatus* chromatophore membranes were added to one mL of assay buffer (25 mM Tris-HCl, pH 7.0, 150 mM NaCl), and the enzymatic reaction was initiated by addition of TMPD at a final concentration of 1 mM. The TMPD oxidation specifically due to *cbb*_3_-Cox activity was controlled by incubating the chromatophore membranes with 200 μM of KCN for two min prior to TMPD addition. The *cbb*_3_-Cox activity was calculated by subtracting from the TMPD oxidase activity the fraction that was KCN-insensitive (15).

### Radioactive ^64^Cu uptake assays

Cellular Cu uptake assays were performed according to (12), using whole cells and radioactive ^64^Cu (1.84 × 10^4^ mCi/μmol specific activity) obtained from Mallinckrodt Institute of Radiology, Washington University Medical School. The. *E. coli* strains harboring appropriate pBAD/Myc-His derivatives with *L*-ara inducible *ccoA* wild type and mutant variants (**Table S1**) were grown overnight in 10 mL of LB medium supplemented with 0.5 % *L*-ara and appropriate antibiotics. Cells were pelleted, washed with the assay buffer (50 mM sodium citrate, pH = 6.5 and 5% glucose) and re-suspended in one mL of the same buffer. All cultures were normalized to the same number of cells (7.5 × 10^8^ cells / 500 μL of assay volume) based on their absorbance (1 OD_600_ = 5×10^8^ cells/mL for *E. coli* and 1 OD_630_= 7.5×10^8^ cells/mL for *R. capsulatus* strains). Cells to be assayed for ^64^Cu uptake were pre-incubated at 35 °C or 0 °C for 10 min before the assay. The uptake activity was initiated by addition of 10^7^ cpm of ^64^Cu, determined immediately before use (half-life of ^64^Cu isotope ~ 12 hrs), and 50 μL of aliquots were taken at 0, 1, 2, 5, and 10 min of incubation time, and immediately mixed with ice-cold 50 μL of 1 mM CuCl_2_ and 50 μL of 50 mM EDTA (pH 6.5) to stop the uptake reaction. All aliquots were kept on ice until the end of the assay, at which time cells were pelleted, pellets washed twice with 100 μL of ice-cold 50 mM EDTA solution, re-suspended in one mL of scintillation liquid, and counted using a scintillation counter (Coulter-Beckman Inc.) with a wide-open window. For each time point, the background ^64^Cu uptake activity seen at 0 °C was subtracted from that at 35 °C and plotted in function of time to compare CcoA-specific Cu uptake of wild type control (*ΔccoA*/plasmid-born *ccoA*) and mutants derivatives of CcoA.

## Abbreviations

Cu: copper
*cbb*_3_-Cox: *cbb*_3_-type cytochrome *c* oxidase
TMH: transmembrane helix
MFS: major facilitator superfamily
CalT: CcoA-like transporters
PMSF: phenylmethylsulfonylfluoride
DTT: dithiothreitol
TCA: trichloroacetic acid
SDS: sodium dodecylsulfate
mPEG-Mal: methoxypolyethylene glycol maleimide
DMPD, *N, N, N’, N’*: dimethyl-*p*-phenylene diamine
TMPD: tetramethyl-*p*-phenylenediamine
N-ter: N-terminal
C-ter: C-terminal
3 D: three dimensional

## Supplemental Materials

Supplemental material for this article may be found at https://

**Table S1.** Strains and plasmids

**Table S2**. Primers used in this study

**Table S3**. Approximate distances separating the Cys residues αC-αC of CcoA

**Figure S1**. CcoA of Rhodobacteriales

**Figure S2**. CcoA of Proteobacteria

## Acknowledgments

This work was supported mainly by the Division of Chemical Sciences, Geosciences and Biosciences, Office of Basic Energy Sciences of Department of Energy [DOE DE-FG02-91ER20052] to FD, partly by the National Institute of Health [NIH GM 38237] to FD, and by the Deutsche Forschungs Gemeinschaft (DFG) (RTG 2202, Project-ID 278002225) to HGK.

## Author Contributions

All authors have given approval to the final version of the manuscript. BK-H, P-IT, and FD designed, performed experiments, and analyzed data. SS did structural analyses, and all authors critically read and edited the manuscript. BK-H, HGK and FD supervised the study.

